# Consideration of sex as a biological variable in diabetes research across twenty years

**DOI:** 10.1101/2023.06.13.544882

**Authors:** Celena Cherian, Hayley Reeves, Duneesha De Silva, Serena Tsao, Katie E. Marshall, Elizabeth J. Rideout

## Abstract

Sex differences exist in the risk of developing both type 1 and type 2 diabetes, and in the risk of developing diabetes-associated complications. Sex differences in glucose homeostasis, islet and β cell biology, and peripheral insulin sensitivity have also been reported in multiple animals. To determine the degree to which biological sex has been addressed in published literature related to diabetes and insulin biology, we developed a scoring system to assess the inclusion of biological sex in papers related to these topics. We scored manuscripts published in *Diabetes*, published by the American Diabetes Association, as this journal focuses on diabetes and diabetes-related research. We scored papers published across three years within a 20-year period (1999, 2009, 2019), a timeframe that spans the introduction of funding agency and journal policies to improve the consideration of biological sex as a variable. Our analysis shows fewer than 15% of papers used sex-based analysis in even one figure across all study years, a trend that was reproduced across journal-defined categories of diabetes research (*e.g*., islet studies, signal transduction). Single-sex studies accounted for approximately 40% of all manuscripts, of which >87% used male subjects only. While we observed a modest increase in the overall inclusion of sex as a biological variable during our study period, our results highlight significant opportunities to improve consideration of sex as a biological variable in diabetes research. In particular, we show that journal policies represent one way to promote better consideration of biological sex as a variable. In the long term, improved practices will reveal sex-specific mechanisms underlying diabetes risk and complications, generating insights to support the development of sex-informed prevention and treatment strategies.

## INTRODUCTION

Diabetes mellitus is a disease that affects millions of individuals globally. According to the International Diabetes Federation, approximately 537 million individuals (20-79 years) worldwide live with diabetes and it has caused 6.7 million deaths^1^. There are also rare monogenic forms of diabetes that affect many people^2^. While diverse factors influence the risk of developing diabetes^3–9^, there is growing recognition that biological sex impacts the risk of developing diabetes and diabetes-related complications^8–13^. In several populations, the risk of developing type 2 diabetes (T2D) is approximately 40% higher in males than in females^8, 9, 11, 12, 14, 15^. In adults aged 15-40, males also have a 60% higher risk of developing type 1 diabetes (T1D)^16^. This male bias in T1D is particularly notable given that females are at a higher risk of nearly all other autoimmune-related diseases^17–19^. After diabetes onset, females are at higher risk of diabetic heart disease, stroke mortality, depression, and anxiety, whereas males show faster progression of nephropathy and require lower extremity amputation more often^9, 20–25^.

Clues into factors that contribute to sex differences in the risk of developing diabetes and diabetes-related complications include a male-female difference in peripheral insulin sensitivity. In humans and many animal models^26–29^, females typically show higher insulin sensitivity^30–34^. Given the established links between reduced peripheral insulin sensitivity and T2D^27, 34–41^, and between insulin sensitivity and diabetes-associated complications^9, 21–25, 42^, biological sex is an important variable to consider when studying insulin sensitivity across physiological and pathological contexts. Sex differences also exist across many animals, including humans, in the biology of insulin-producing β cells^30, 43–47^. Male-female differences have been reported in the number of pancreatic β cells^47^, and studies in humans and rodents report profound sex differences in β cell gene expression, function, and stress responses in both normal and T2D contexts^30–34, 48, 49^. Female β cells also show higher glucose-stimulated insulin secretion under normal physiological conditions and in T2D^9, 30–32^, differences that cannot be solely attributed to sex differences in peripheral insulin sensitivity^30^. Because there are established links between β cell dysfunction in rodent models and humans in T2D^49–55^, and emerging evidence suggests β cell dysfunction may also contribute to T1D^56, 57^, biological sex is an important variable to consider in studies on islets and β cells in multiple physiological and pathological contexts. Yet, the degree to which biological sex has been considered in the literature with respect to islet and β cells is incompletely known.

Broad-based text searches of diabetes articles, and detailed assessments of randomized controlled trials and human observational studies in top diabetes journals, suggest that biological sex is rarely considered as a variable in diabetes research^58–61^. Indeed, one study showed that only 6.5% of randomized controlled trials and human observational studies reported all outcomes separately according to biological sex^58^. While these studies were critical in highlighting the overall poor consideration of both sexes in diabetes research, several questions remain. For example, the prior focus on randomized controlled trials and observational studies means that we lack detailed information on whether biological sex is included in studies that focus on cell, tissue, and animal models of diabetes. We similarly lack information on whether there are differences in practices related to biological sex between different diabetes subject areas (*e.g*. islet biology, transplantation), and whether funding agency policies mandating the inclusion of both sexes^59, 62–65^ lead to better consideration of biological sex in diabetes research. Answering these questions will highlight opportunities for improvement in research design, methods, and analysis across all areas of diabetes research, and establish whether policies introduced by major funding agencies, journals, and governments have produced sufficient change in research practices^62–66^.

We performed a detailed analysis of all manuscripts published in the journal *Diabetes*, published by the American Diabetes Association, in 1999, 2009, and 2019 (819 publications) (Fig. 1A). Studying the inclusion of sex as a biological variable in one journal rather than between journals provides advantages due to uniform journal policies and similar manuscript quality. Because *Diabetes* publishes papers in multiple subject areas, and in both clinical and biomedical disciplines, we captured and compared differences in common practices across diverse fields related to diabetes mechanisms and risk. Overall, our findings suggest that fewer than 15% of studies published in *Diabetes* during our study period analyzed data using biological sex as a variable. Approximately 34% of studies included both sexes in data collection, but did not analyze their data using biological sex as a variable, and 40% of studies included only a single sex. While we observed a significant increase in the inclusion of both sexes in diabetes research over time, we observed a smaller increase in the number of studies that used sex-based analysis over time. These trends were broadly similar across many subfields of diabetes research, and funding status had no significant effect on overall consideration of biological sex as a variable in diabetes research. Together, our data suggests progress has been made in diabetes-related research with respect to the inclusion of both sexes and uptake of sex-based analysis; however, significant opportunities for improvement remain. This is especially true for single-sex studies, the vast majority of which used male and not female animals. Greater knowledge of how biological sex affects diabetes mechanisms and complications will ultimately support the development of sex-informed prevention and treatment strategies to improve equity in health outcomes^61, 66^.

**Figure 1.**
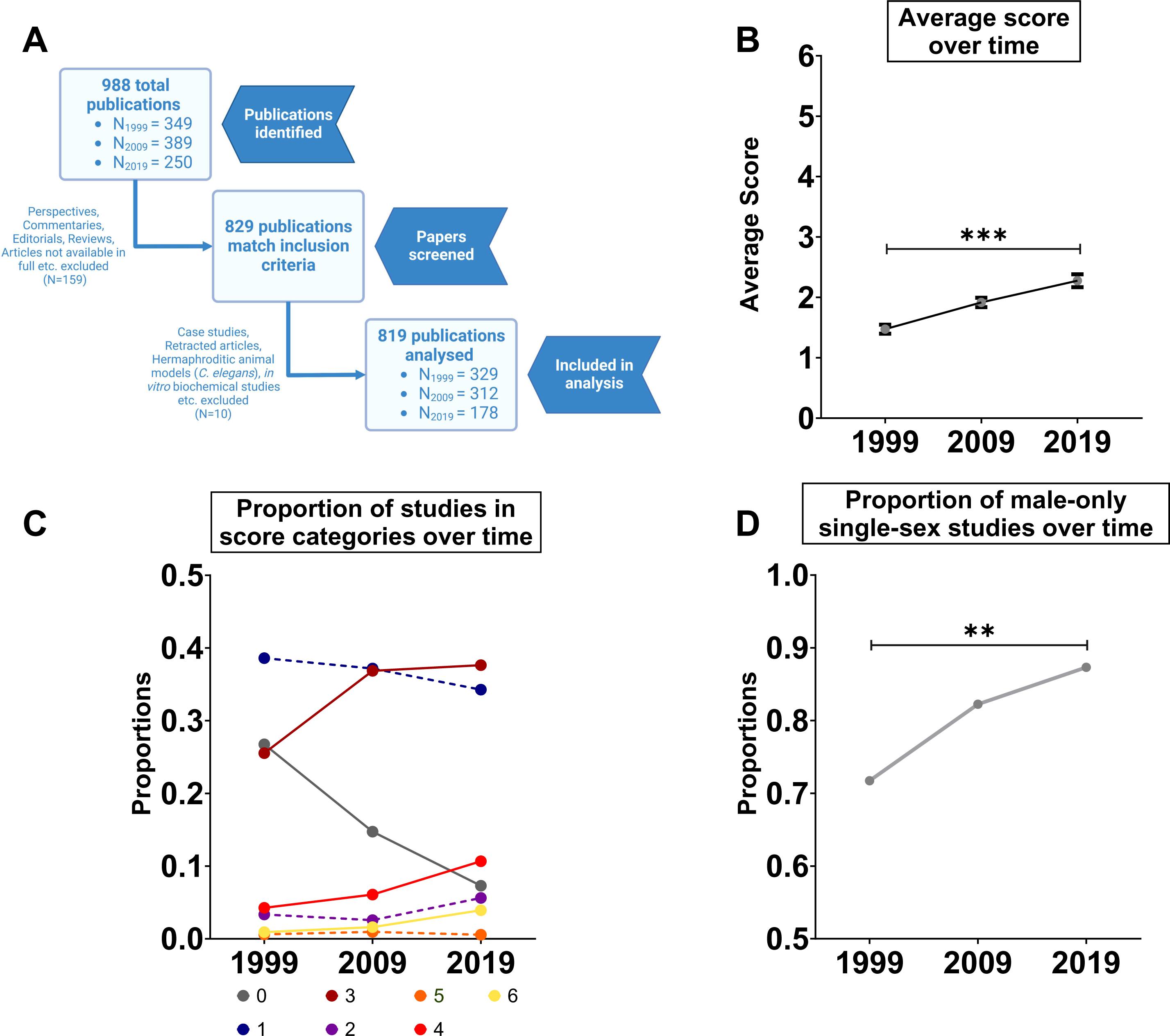
Assessing the inclusion of biological sex as a variable in *Diabetes* manuscripts published over time. (A) Schematic representation of the workflow for manuscript selection in our study. (B) Average score of studies published in 1999, 2009, and 2019. (*** p<0.001; generalized linear model with Poisson distribution; error bars indicate SEM) (C) Proportion of studies assigned scores between 0-6 in 1999, 2009, and 2019. Dashed lines represent a statistically insignificant change and solid lines represent a statistically significant change (generalized linear model with binomial distribution). (D) Proportion of single-sex studies that used male animals in 1999, 2009, and 2019 (** p<0.01; generalized linear model with binomial distribution).

## RESULTS

### Assessing the inclusion of biological sex as a variable in diabetes research over time

The start of the twenty-year period we considered in our study corresponded with the introduction of major policy initiatives aimed at improving the consideration of biological sex as a variable in Canada, Europe, and the United States^62–66^. To determine whether the inclusion of biological sex as a variable in diabetes research changed over time, we monitored the average score of the papers in our study in 1999, 2009, and 2019. The average score of papers published in 1999 was 1.47 ± 0.07, whereas the average scores of manuscripts published in 2009 and 2019 were 1.91 ± 0.073 and 2.28 ± 0.1, respectively (Fig. 1B). While this represents a significant increase in average score between 1999 and 2019 (year effect *p*<0.001; ANOVA in a generalized linear model with Poisson distribution), an increase in average score of only 0.8 over 20 years indicates that significant room for improvement remains in the inclusion of sex as a biological variable in diabetes research.

To describe the dynamics underlying the change in average score over time in more detail, we examined the proportion of studies classified into each of our six categories. The proportion of studies assigned a score of 0 significantly declined over time, from 27% in 1999, to 19% in 2009, and 7.3% in 2019 (year effect *p*<0.001; ANOVA in a generalized linear model with binomial distribution) (Fig. 1C). In contrast, there was a significant increase in the proportion of papers over time assigned a score of 3, 4, or 6 (year effect *p*=0.002, *p*=0.004, *p*=0.029 respectively; ANOVA in a generalized linear model with binomial distribution) (Fig. 1C). Papers assigned a score of 3 comprised 25.5% of papers in 1999, 36.8% in 2009, and 37.6% in 2019 (Fig. 1C). Papers assigned a score of 4 made up 4.2% of papers in 1999, 6% papers in 2009, and 10.6% papers in 2019, whereas papers assigned a score of 6 represented only 0.91% of studies in 1999, 1.6% in 2009, and 3.9% in 2019 (Fig. 1C).

No significant change was observed in the proportion of papers scored as a 1, 2, or 5 between 1999 and 2019 (year effect *p*=0.618, *p*=0.519, *p*=0.991 respectively; ANOVA in a generalized linear model with binomial distribution) (Fig. 1C). Based on this data, the categories with the most pronounced changes between 1999 and 2019 were 0 (decrease) and 3 (increase) (Fig. 1C). This suggests the higher average score in 2019 may be partly attributed to a drop in studies that did not discuss biological sex at all and an increase in mixed-sex studies with no sex-based analysis. Increased inclusion of both sexes represents a promising step toward better consideration of biological sex as a variable in diabetes research; however, important next steps for improvement include greater use of sex-based analysis.

### Male animals are used more often in single-sex studies

Previous studies on the inclusion of biological sex as a variable in biomedical research reveal that male animals are used more frequently than female animals^59, 60, 67^. For example, a study from 2010 analyzed almost 2,000 animal studies across 10 biological fields in several journals and found a male bias in eight fields, with the most pronounced bias in neuroscience, pharmacology, and physiology^60^. To determine whether one sex was used more often in diabetes research, for all papers assigned a score of 1 or 2 (single-sex studies) we recorded the sex that was used. Studies assigned a score of 1-2 represented 42% of studies in 1999, 39.7% of studies in 2009, and 40% of studies in 2019 (Fig. 1D), suggesting single-sex studies represented a large proportion of total studies published during this period. In 1999, 71.7% of single-sex studies used only male animals (Fig. 1D). In 2009 and 2019, 82.2% and 87.3% of single-sex studies used only male animals, respectively (Fig. 1D). When we examined the number of male-only studies over time, we observed a significant increase between 1999 and 2019 (year effect *p*=0.023; ANOVA in a generalized linear model with binomial distribution) (Fig. 1D). This suggests that in diabetes research, as in other fields of biomedical science^59, 60, 67^, male animals are used more frequently than female animals.

### Evaluating the inclusion of biological sex as a variable in diabetes research across clinical and biomedical disciplines

Diabetes research includes both clinical and biomedical disciplines. Given that policies governing the inclusion of biological sex as a variable were introduced at an earlier date for clinical research compared with biomedical research^59, 60, 68, 69^, we compared the average score assigned to studies in each of these disciplines in 1999, 2009, and 2019. Across all three study years, the average score for papers in the clinical discipline (2.78 ± 0.08) was significantly higher than the average score for papers in the biomedical discipline (1.48 ± 0.05) (discipline effect *p*<0.001; ANOVA in a generalized linear model with Poisson distribution) (Fig. 2A). We observed a significant increase in the average scores across both biomedical and clinical disciplines between 1999 and 2019 (year: discipline interaction *p*<0.0017; ANOVA in a generalized linear model with Poisson distribution) (Fig. 2A), with a trend toward a larger increase over time in the biomedical discipline.

**Figure 2.**
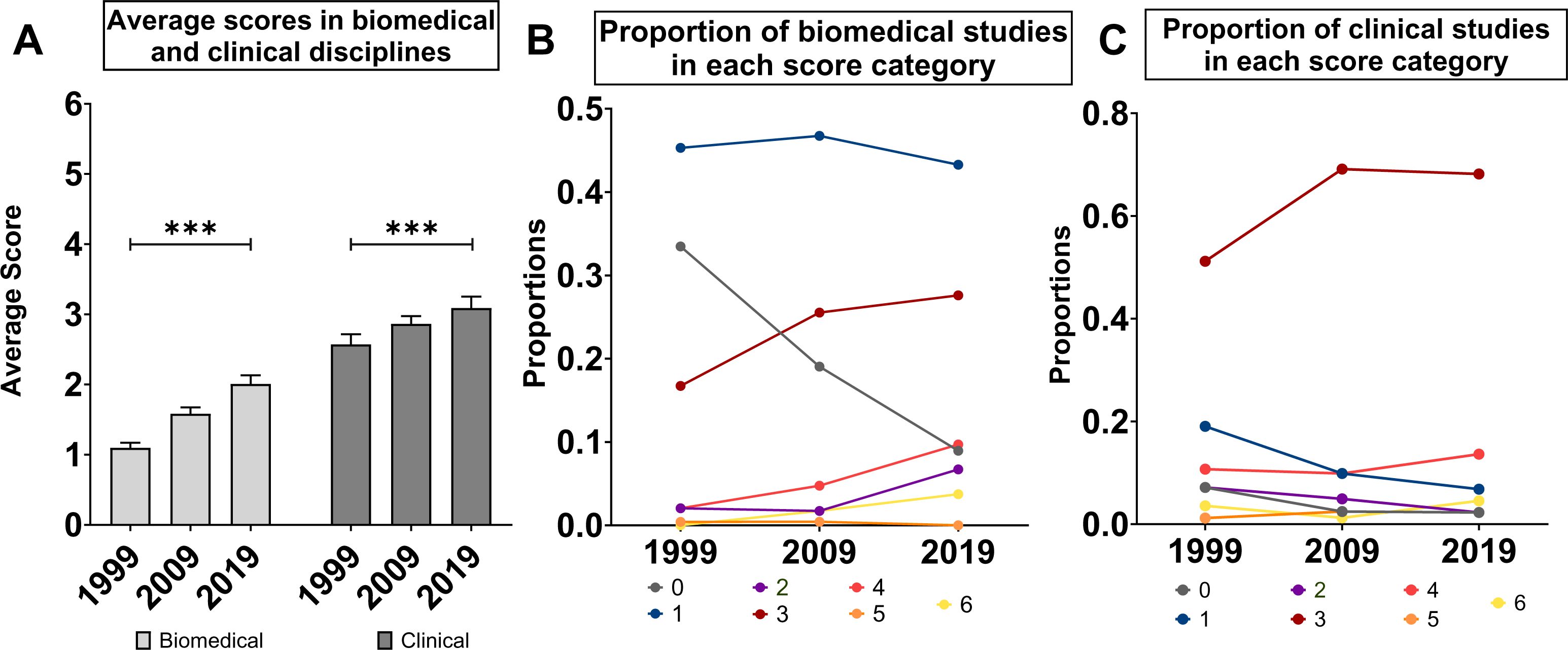
Evaluating the inclusion of biological sex as a variable in *Diabetes* manuscripts across clinical and biomedical disciplines. (A) Average scores assigned to studies published in biomedical and clinical disciplines in 1999, 2009, and 2019 (*** p<0.001; generalized linear model with Poisson distribution; error bars indicate SEM). (B) Proportion of biomedical studies assigned scores between 0-6 in 1999, 2009, and 2019. (C) Proportion of clinical studies assigned scores between 0-6 in 1999, 2009, and 2019.

To identify factors that contribute to the score difference between the biomedical and clinical disciplines, we examined the probability that a study in each category would be assigned a specific score (Fig. 2B,C). While there was a significant decrease in the proportion of both biomedical and clinical studies assigned a score of 0 over time (year effect *p*<0.001; ANOVA in a generalized linear model with binomial distribution), the probability that a biomedical study would be assigned a score of 0 was higher than the probability of a clinical study being assigned this score across all study years (discipline effect *p*<0.001; generalized linear model with binomial distribution) (Fig. 2B,C). The probability of a biomedical study receiving a score of 1 was also significantly higher than the probability of a clinical study being assigned this score in all three study years (discipline effect *p*<0.001; generalized linear model with binomial distribution) (Fig. 2B).

In the clinical discipline, the score that was assigned to the highest proportion of studies across all years was 3 (61.7% of studies; Fig. 2C). Indeed, the probability of a clinical study being assigned a score of 3, 4 or 5 was significantly higher than the probability of a biomedical study being assigned these scores in all study years (discipline effect *p*<0.001, p=0.002 and *p*=0.03, respectively; generalized linear model with binomial distribution) (Fig. 2B,C). While our data indicate that more authors indicated the sex of the model system they used in 2019 than in 1999, significant room for improvement remains. For example, in 2019 the majority of biomedical studies still used only a single sex (∼50%) and most failed to use sex-based analysis (86.6%). In the clinical discipline, fewer than 15% of published studies used sex-based analysis during our study period.

### Comparing the inclusion of biological sex as a variable between diabetes subject areas

Beyond the diverse approaches to diabetes research between clinical and biomedical disciplines, diabetes research spans a broad range of topics, from genetic and genomic approaches, to immunology, to pharmacology. We therefore wanted to compare the inclusion of biological sex as a variable across different diabetes subject areas. From 2009 onwards, all studies published in *Diabetes* correspond to one of 10 primary topics/subjects (Table 2). Note that not all areas maintained identical labels in 2009 and 2019. For example, the topic ‘Genetics’ in 2009 was compared to ‘Genetics/Genomes/Proteomics/Metabolomics’ in 2019 (hereafter called Genetics/Omics). Similarly, the ‘New Methodologies and Databases’ was compared to ‘Technological Advances’ in 2019 (hereafter called New Methodology and Technology). To ensure adequate sample sizes within each topic, we excluded subject areas with fewer than ten papers in either 2009 or 2019 (Signal Transduction, New Methodology and Technology, Pharmacology & Therapeutics).

In 2009, the average score was highest in the Genetics/Omics and Obesity subject areas (2.5 ± 0.17 and 2.62 ± 0.39, respectively), and the lowest in Immunology and Transplantation and Islet topics (1.62 ± 0.27 and 1.35 ± 0.23, respectively; Fig. 3A). In 2019, the average scores were highest in the Genetics/Omics, Obesity, and Immunology and Transplantation topics (3.04 ± 0.20, 2.67 ± 0.34, and 2.72 ± 0.38, respectively) (Fig. 3A). In both 2009 and 2019, the higher score for studies under the Genetics/Omics subject area may be attributed to the higher probability of a study in that subject area being assigned a score of 3 compared with other subject areas (Fig. 3B-H). Despite the different average scores between subject areas, our model suggests that all subject areas showed a similar change in average score over time (year:subject interaction *p*=0.341; ANOVA in a generalized linear model with Poisson distribution). Thus, our analysis did not reveal widespread differences in practices related to the inclusion of biological sex as a variable between different diabetes subject areas.

**Figure 3.**
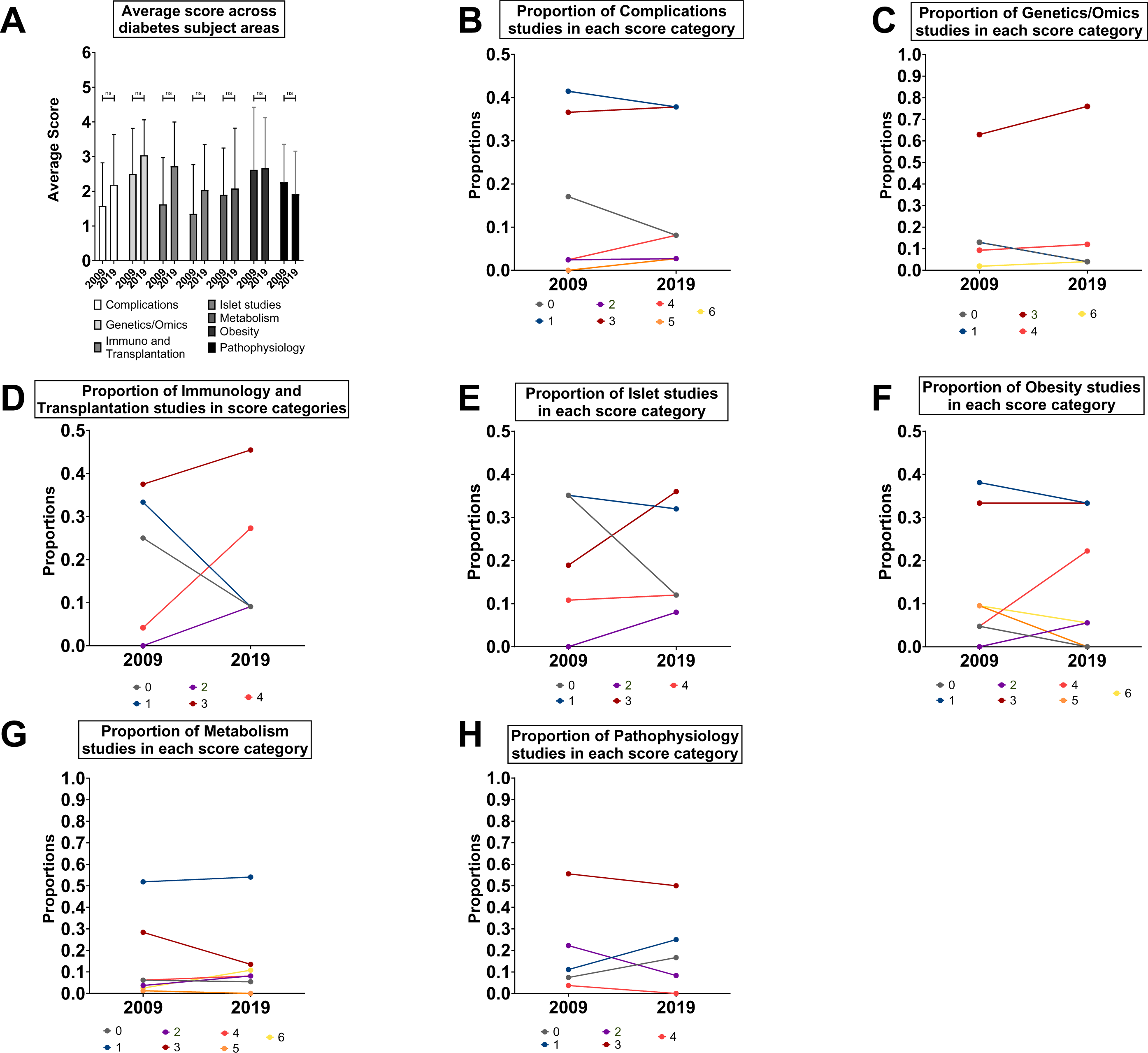
Comparing the inclusion of biological sex as a variable between *Diabetes* manuscripts published across diabetes subject areas. (A) Average scores assigned to studies published in different diabetes subject areas in 2009 and 2019. (ns not significant; generalized linear model with Poisson distribution; error bars indicate SEM) (B) Proportion of studies assigned scores between 0-6 in 2009 and 2019 in the ‘Complications’ subject area. (C) Proportion of studies assigned scores between 0-6 in 2009 and 2019 in the ‘Genetics/Omics’ subject area. No studies in either year were assigned a score of 2 or 5. (D) Proportion of studies assigned scores between 0-6 in 2009 and 2019 in the ‘Immunology and Transplantation’ subject area. No studies in either year were assigned a score of 5 or 6. (E) Proportion of studies assigned scores between 0-6 in 2009 and 2019 in the ‘Islet’ subject area. No studies in either year were assigned a score of 5 or 6. (F) Proportion of studies assigned scores between 0-6 in 2009 and 2019 in the Obesity subject area. (G) Proportion of studies assigned scores between 0-6 in 2009 and 2019 in the Metabolism subject area. (H) Proportion of studies assigned scores between 0-6 in 2009 and 2019 in the Pathophysiology subject area. No studies in either year were assigned a score of 5 or 6.

### Evaluating the effect of funding source on the inclusion of biological sex as a variable in diabetes research

Funding agencies across many countries have introduced policies aimed at increasing the inclusion of biological sex as a variable in research^59, 62–66^. The US National Institutes of Health (NIH) represents a key source of funds for health research, with significant investments in diabetes research^63^. The NIH was among the first funding agencies worldwide to introduce policies related to the inclusion of both sexes in both clinical and biomedical research^59, 62, 64–66^. To evaluate whether NIH funding affected the inclusion of biological sex as a variable in diabetes research, we compared the average score assigned to studies by authors that declared NIH funding with scores of studies by authors that did not declare NIH funding (Fig. 4A). We found that the average score of NIH-funded studies was not significantly higher than studies that declared funding from other sources (funding effect *p*=0.067; year:funding interaction p=0.974; ANOVA in a generalized linear model with Poisson distribution) (Fig. 4A). We next compared the proportion of studies within each score category between NIH-funded studies and non-NIH-funded studies (Fig 4B,C). For the majority of score categories (0-5), NIH funding had no significant effect on the proportion of studies assigned a specific score (funding effect *p*=0.942, p=0.236, p=0.777, p=0.589, p=0.771, p=0.222, respectively; ANOVA in a generalized linear model with binomial distribution) (Fig. 4B,C). In contrast, NIH-funded studies showed a significantly higher proportion of studies assigned a score of 6 across all study years compared with non-NIH-funded studies (funding effect *p*<0.001; funding:year interaction *p*=0.049; ANOVA in a generalized linear model with binomial distribution) (Fig. 4B,C). This data indicates that NIH funding was not associated with overall shifts in practices related to the consideration of biological sex as a variable in diabetes research. Indeed, despite the fact that policies related to the inclusion of biological sex as a variable in clinical research^16^ were introduced earlier than similar policies related to biomedical research^62, 64, 65^, there was no significant interaction between year, funding source and type of study when we asked whether NIH funding affected the average scores assigned to clinical and biomedical studies (Fig. 4D). This reinforces our conclusion that NIH funding was not associated with increased consideration of biological sex as a variable in diabetes research.

**Figure 4.**
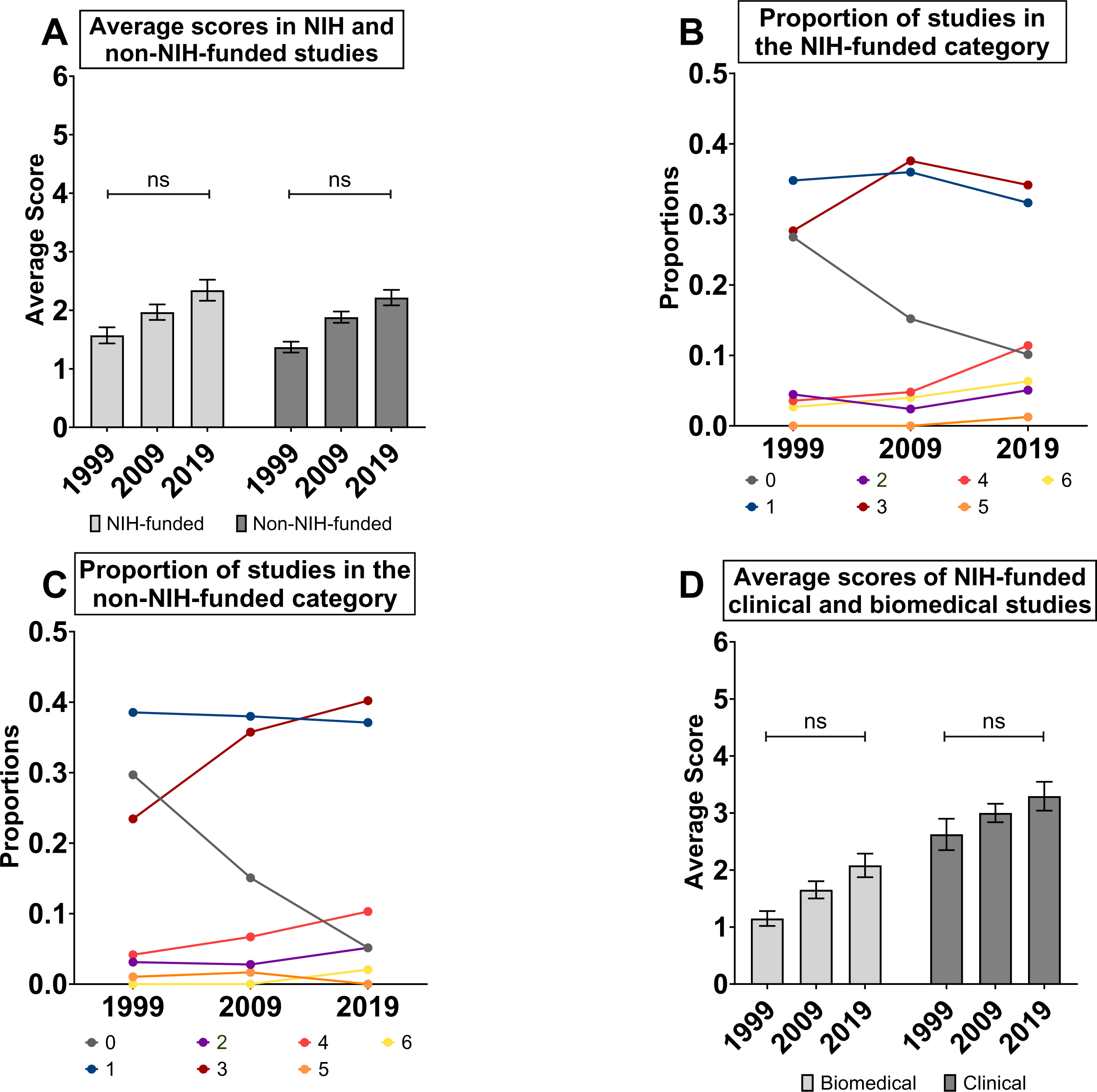
Evaluating the effect of funding source on the inclusion of biological sex as a variable in diabetes research. (A) Average scores assigned to studies that declared NIH funding compared with studies that did not declare NIH funding in 1999, 2009, and 2019 (ns not significant; generalized linear model with Poisson distribution; error bars indicate SEM). (B) Proportion of NIH-funded studies assigned scores between 0-6 in 1999, 2009, and 2019. (C) Proportion of non-NIH-funded studies assigned scores between 0-6 in 1999, 2009, and 2019. (D) Scores assigned to studies that declared NIH funding in the biomedical and clinical disciplines in 1999, 2009, and 2019 (ns not significant; generalized linear model with Poisson distribution; error bars indicate SEM).

### Evaluating the effect of journal policies on inclusion of sex as a biological variable

In 2019, a new policy was introduced at *Diabetes* mandating that authors of studies that use human islets provide detailed information on donor characteristics^70–72^. The goal of including this detailed donor information was to enhance transparency and to enable better comparisons among data generated using human islets. One of the required donor characteristics included in the Human Islet checklist is biological sex^72^. Because disclosing the sex of experimental models represents one way that authors can improve consideration of biological sex in their research, we assessed the effect of this policy change on the scores assigned to papers in the ‘Islet studies’ subject area before and after the policy came into effect in April 2019. We scored all manuscripts published in this subject area in 2018 (36 papers) and 2020 (32 papers), as this was the topic most likely to be affected by a policy regarding human islet documentation. We did not score papers published in 2019 as these papers were likely submitted and/or reviewed prior to the policy change.

We found the average score of papers in the Islet subject area in 2018 was 1.66 ± 0.64, whereas the average score of studies in this area was 2.5 ± 0.19 in 2020 (Fig. 5A). This represented an increase of 0.8 in the average score of studies in the Islet subject area over only a two-year period (Fig. 5A). While this increase was not statistically significant (year:subject interaction *p*=0.248; ANOVA in a generalized linear model with Poisson distribution) it is worth noting that the magnitude of this increase is equivalent to the increased we observed in all studies across our twenty-year study period. Potential explanations for the increase in average score for studies in the Islet subject area include a trend toward more studies assigned a score of 4-6, and a trend toward fewer studies assigned a score of 0-1 (Fig. 5B). To determine whether this increase was reproduced in another subject area less likely to be affected by the policy change regarding human islets, we repeated our analysis of manuscripts published in 2018 and 2020 in the ‘Metabolism’ subject area. The average score for studies in the ‘Metabolism’ subject area increased by 0.03 over the span of the policy change and was not significantly different in 2018 compared with 2020 (year:subject interaction *p*=0.248; ANOVA in a generalized linear model with Poisson distribution) (Fig. 5A). Taken together our analysis suggests that the editorial policy change at *Diabetes* may have contributed to a significant improvement in the consideration of biological sex as a variable in diabetes research.

**Figure 5.**
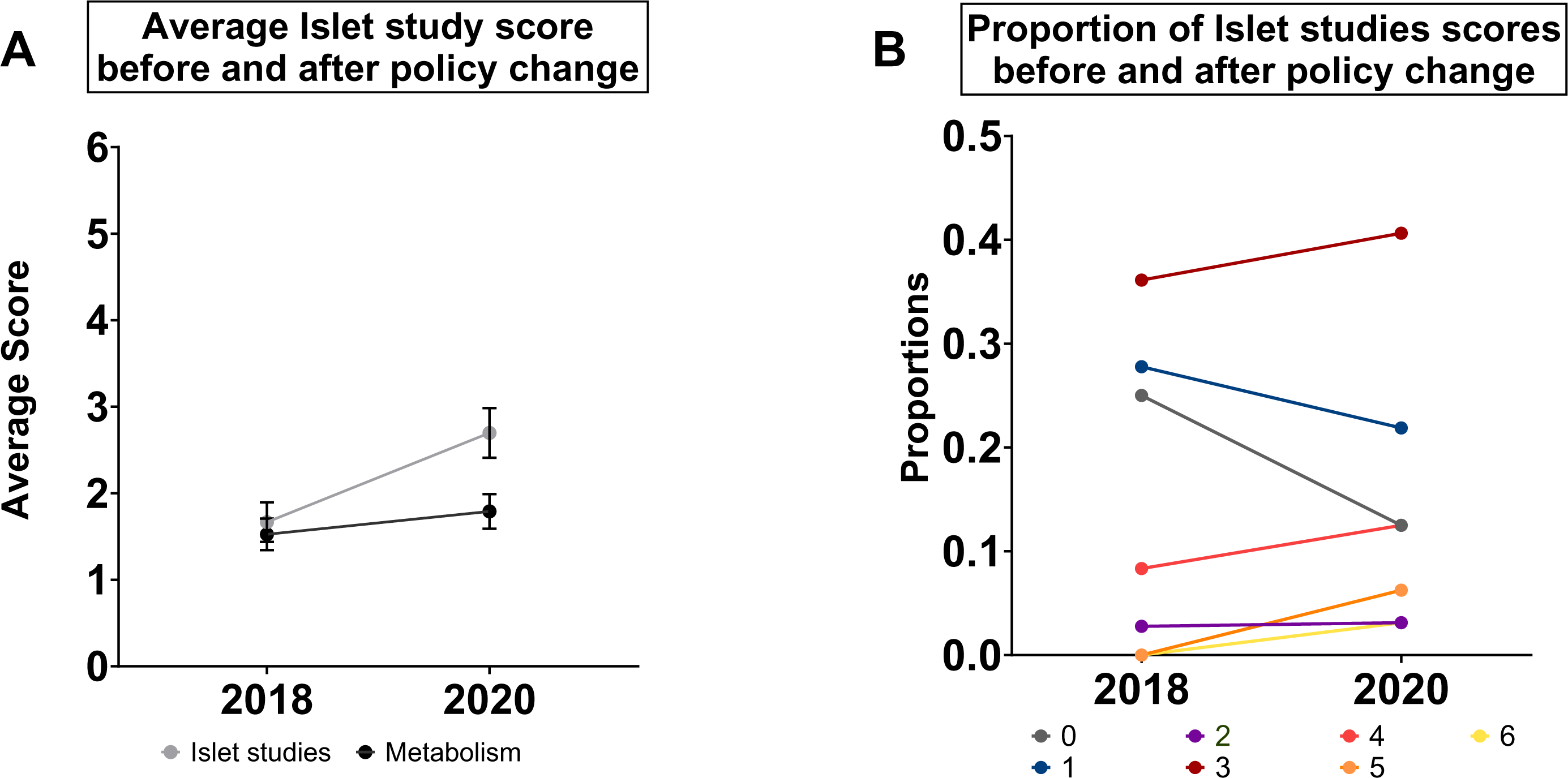
Evaluating the effect of journal policies on inclusion of sex as a biological variable in *Diabetes* manuscripts. (A) Average scores assigned to studies in the ‘Islet’ and ‘Metabolism’ subject areas between 2018 and 2020 (before and after the 2019 policy change, respectively) (** p<0.01; ns not significant; generalized linear model with Poisson distribution; error bars indicate SEM). (B) Proportion of studies in the ‘Islet’ subject area assigned scores between 0-6 in 2018 and 2020.

## DISCUSSION

The overall goal of our study was to assess the degree to which biological sex was included as a variable in diabetes research across a twenty-year period. We chose to assess studies in a journal that is dedicated to publishing high-quality studies in the area of diabetes research, and across a timeframe that captures widespread changes to journal and funding agency policies regarding sex- and gender-based analysis^62, 64, 65, 70–72^. While our analysis revealed significant increases in the inclusion of biological sex as a variable over time, in alignment with the reported increase in the word “sex” in text-based searches of a similar body of literature^58^, our detailed evaluation of methods demonstrates that many studies do not adequately address biological sex as a variable in diabetes research. Given that biological sex affects diabetes risk^1, 9, 16, 56^, treatment efficacy^3, 8, 71^, and risk of developing diabetes complications^9, 20, 21, 25, 42^, this knowledge gap must be closed in order to develop sex-informed prevention and treatment strategies for people living with diabetes.

One major barrier to the full consideration of biological sex as a variable in diabetes research was related to inclusion. In all three study years across a twenty-year period, between 46 and 69% of studies did not include both sexes (score 0-2). This low inclusion of both sexes was particularly notable in biomedical studies (between 59% and 81% of studies across the 3 years were assigned a score of 0-2) and in diabetes subject areas that contained a large proportion of biomedical studies (*e.g.,* islets and metabolism). A lack of inclusion of both sexes has been described in studies of literature related to neuroscience, psychiatry and psychology, pharmacology, and physiology^60, 67, 68, 73, 74^, and our data aligns with findings from others^58, 68, 69, 73, 75, 76^ suggesting similar trends exist in diabetes research. Indeed, the inclusion of both sexes did not show a meaningful increase over a twenty-year period in many subject areas of diabetes research. Addressing this overall lack of inclusion is an important task going forward for the diabetes community as funding agencies and journals worldwide aim to improve transparency and reproducibility in research^59, 66^. Because our study showed the majority of single-sex studies use only male subjects, the major change that is needed in biomedical studies is the inclusion of female subjects. Indeed, our analysis suggests that the gap in knowledge of female cells and animals will not be closed if current trends in single-sex studies continue. Given that female subjects are protected from dysregulation of glucose homeostasis^27, 30–32, 34, 37, 38^, but are at a higher risk of developing complications from diabetes^9, 21, 42^, mechanistic insights gained from studying females will provide valuable insights into diabetes prevention and treatment.

While a lack of inclusion of both sexes is an important barrier to consideration of biological sex as a variable in diabetes research, it is important to note that single-sex studies are acceptable in some cases. For this reason, we assigned a higher score (2) to a small number of manuscripts (n=34) that provided a reason for their use of a single-sex study group. When we examined the reasons provided by the authors of manuscripts assigned a score of 2 for their choice of using a single-sex study group, we found two broad categories of justifications. In the first category, the reason was that the single-sex study group included pregnant people or individuals living with polycystic ovary syndrome. Because these contexts are not experienced by both sexes, it is acceptable to use a single-sex group in these studies. Future studies will need to adjust our scoring system to reflect the fact that using single-sex study groups with a strong justification considers biological sex properly, and should perhaps score higher than mixed-sex studies.

The second category of studies assigned a score of 2 included manuscripts that provided a variety of reasons for including only a single sex. Reasons given for using a single-sex study group in this category included a lack of prior knowledge in one sex and fluctuating female hormones (Supplemental file 1). While a lack of detailed knowledge of diabetes mechanisms in both sexes is an unfortunate consequence of the fact that few studies properly consider biological sex as a variable, it does not represent a strong justification for using a single-sex study group. Similarly, several studies now show that female animals are not more variable than males across multiple traits^77, 78^. These are therefore not strong justifications for the failure to include both sexes in a study.

Another major barrier to the full consideration of biological sex as a variable in diabetes research was a lack of sex-based analysis. Studies assigned a score of 3 comprised between 17% and 28% of those published in the biomedical discipline and between 51% and 77% of clinical studies between 1999 and 2019. In the biomedical discipline, the most common reason studies were assigned a score of 3 was that they pooled samples (*e.g*., islets) collected from male and female experimental models prior to collecting measurements. This suggests that in biomedical studies, the major changes that are needed to support sex-based analysis are to 1) collect samples and data from male and female experimental models separately, and 2) to use appropriate statistical methods to analyze the data from each sex separately. In clinical studies, the main reason studies were assigned a score of 3 was that biological sex was not treated as a variable. Instead, most studies in the clinical discipline considered biological sex as a covariate. Given that biological sex can interact with treatment to drive outcomes, including biological sex only as a covariate can mask sex-dependent effects. Biological sex should therefore be included either as an interactor, or analyses should be stratified by sex^74^. The main change that is needed in the clinical discipline is therefore to adjust study analysis methods to fully consider biological sex as a variable. Because many studies show sex differences in islet and β cell function^41, 47, 48, 79^, the genetic architecture of disease^9, 41, 42, 55^, and metabolic differences^27, 30, 32, 33, 38^, shifts in practice across the biomedical and clinical disciplines, and in different diabetes subject areas, will allow researchers to develop evidence-based models for these topics in each sex. These models will form the basis for the development of sex-informed prevention and treatment efforts for individuals at risk for, or living with, diabetes.

Despite the growing number of studies showing that biological sex is not given adequate consideration in biomedical and clinical research^60, 68, 69, 80^, the most effective strategies to promote better uptake of biological sex as a variable are not fully understood. Our data, in alignment with similar studies^73, 75, 76^, indicate that funding by agencies with clearly-stated requirements related to the consideration of biological sex does not significantly influence author practices (Fig. 4). One potential reason for this lack of influence on author practices is that monitoring compliance with sex- and gender-related policies poses multiple challenges. Due to a policy change during the course of our study, we were able to assess how journal policies affect the consideration of biological sex as a variable. Our data suggests that journal policies may represent an effective tool in promoting the uptake and integration of biological sex as a variable in diabetes research. In the ‘Islet studies’ subject area, the increase in average score of 0.8 between 2018 and 2020 was similar in magnitude to the 0.91 increase in average score across the biomedical discipline over a twenty-year period. This suggests the implementation of robust journal policies related to integration of biological sex as a variable will promote better practices in the future. Applying these policies will require a culture shift across editors, reviewers, and authors, however, proper consideration of biological sex as a variable will provide significant benefits in terms of data reproducibility, transparency, and scientific rigor^69, 73, 75, 80^.

Based on our analysis, we suggest several practices authors can adopt to improve consideration of biological sex as a variable in their research. First, we discuss changes that can be made in the short term. For example, one way to immediately improve transparency regarding integration of biological sex as a variable is to ensure the sex of experimental subjects and materials are explicitly stated in the methods section and in figure legends. If only a single sex is used, justification should be provided for the use of a single-sex experimental group. Authors should also identify the use of a single sex as a limitation at some point in the paper, and state that future studies will be needed to determine whether findings made in one sex apply to the other. This is important even in studies that use cell lines or tissues that are limited to one sex (*e.g*., Chinese hamster ovary cell line), as it allows readers to appreciate potential limitations of the mechanisms uncovered by the authors. If both sexes are used, but not all phenotypes are recorded in each sex, the figure legends should include this information to ensure the reader is aware of which sex was used to generate specific data points.

It is also important to ensure data from each sex is collected and analyzed separately, and not pooled. Pooling male and female cells or tissues during analysis risks overlooking important effects if a sex-specific outcome does not achieve statistical significance within a mixed-sex group. Similarly, potential interactions between biological sex and other variables should be explicitly tested for, or the analysis should be stratified by sex, to allow for detection of sex-specific effects. Treatment of biological sex as a covariate potentially masks those effects. Because these suggestions are all related to changes in text, and how data are displayed, described, and analyzed, they can be applied to studies that are either complete or in progress. Longer-term changes include using male and female cell lines and animal models, collecting and analyzing data from males and females separately, and applying statistical methods to reveal how biological sex affects diverse phenotypes. The solid foundation of knowledge that will emerge from these studies will help overcome the current lack of information on male and female cells and animal models. It will also serve as a first step in embracing diversity in sex chromosome complement and sex hormone levels, in line with emerging recognition that biological sex is not a binary variable^13, 61, 69, 74, 76, 80^.

In conclusion, better knowledge of normal physiology in males and females will allow the development of sex-informed prevention efforts, and better knowledge of pathophysiology will support the discovery of more effective treatment options that reduce the burden of complications. Broader consideration of biological sex as a variable in diabetes research will therefore provide significant benefits to all individuals at risk for, and living with, diabetes.

## METHODS

### Article selection criteria

Our analysis included original research articles for which the full text was available on the *Diabetes* online archive. We did not include reviews, editorials, or commentaries in our analysis. We also excluded case studies, as reports on individuals preclude the meaningful analysis of both sexes (Fig. 1A).

### Scoring the inclusion of sex as a biological variable

Each paper was scored based on the degree to which biological sex was integrated into study design and data analysis. A full breakdown of the scoring system is included as Table 1. Briefly, a score of 0 was assigned to studies in which the sex of animal models or participants was not stated at any point in the text including the methods section. This includes studies that used cell lines with an obvious sex (*e.g*., Chinese hamster ovary cell line) in cases where the authors did not discuss the sex of this cell line as a potential study limitation. A score of 1 was assigned to studies that used a single sex (male or female) but did not provide any rationale for this choice. A score of 2 was assigned to studies that used a single sex but provided an explicit reason for this decision. In some cases the authors provided strong justifications for the use of a single sex, whereas other authors provided less strong justifications for their choice. Strong justifications for the use of a single sex included studies on pregnant subjects and individuals with polycystic ovary syndrome. Less strong justifications included a lack of prior studies in one sex. A score of 3 was given to mixed-sex studies. These studies included both sexes, but did not 1) indicate the number of male and female subjects used and 2) analyze their data by sex. Clinical studies that used biological sex as a covariate were also assigned a score of 3 because this type of analysis does not properly consider biological sex effects as a variable. A score of 4 was given to studies that included biological sex as a variable in the analysis for as little as one piece of data (*e.g*., one panel of a figure reported male and female phenotypes separately). A score of 5 was assigned to studies that included biological sex as a variable in >50% of the data presented in the manuscript. A score of 6 was assigned for studies that included both males and females across all experiments, and in which all data was analyzed and reported by sex. The full data set is available in Supplemental Files 1 and 2.

**TABLE 1.** Scoring system to assess the consideration of biological sex as a variable.

| Score | Meaning |
| --- | --- |
| 0 | No mention of sex |
| 1 | One sex used, without explicit reasoning |
| 2 | One sex used, with explicit reasoning; includes studies with strong and less strong justifications for this choice |
| 3 | Both sexes included; data not analyzed separately by sex or data not provided |
| 4 | Both sexes included; fewer than half of results analyzed by sex (<50%) |
| 5 | Both sexes included; more than half of results analyzed by sex (>50%) |
| 6 | Both sexes included; all data analyzed and reported by sex |

### Scoring reproducibility

After the full data set was collected, a subset of articles was re-scored by another person to assess inter-individual scoring variability. Of the 86 papers that were selected at random for re-scoring, 96.5% (83/86) were assigned the same score by different individuals. This indicates consistency between individuals in the application of our scoring system.

### Recording the sex of experimental subject

For papers with a score of ‘1’ or ‘2’ the sex of the experimental subjects, as reported by the authors, was recorded in our spreadsheet.

### Recording diabetes discipline and subfields

Each paper was classified as either ‘clinical’ or ‘biomedical’ to allow a comparison between these two disciplines regarding the inclusion of biological sex as a variable. Clinical studies included randomized controlled trials and studies in which subjected human participants were subjected to treatment protocols. Observational studies involving human subjects were also classed as clinical when material or recordings beyond a single blood sample, superficial tissue, or DNA sample was taken for analysis. All other papers, including those which used animal models or cell lines, were classed as biomedical. For papers with both clinical and biomedical data, the classification was based on the type of data from which the majority (>50%) of the results were generated. For papers published in 2009 and 2019 the research subject area of each study was also recorded based on the classification system used by *Diabetes* (*e.g*., ‘Islet biology’, ‘Obesity’; see Table 2). Because papers published in 1999 were not categorized according to subject area, this information was only collected for 2009 and 2019.

**TABLE 2.** Number of papers in each diabetes subject area.

| Diabetes subject area | Number of papers in 2009 | Number of papers in 2019 |
| --- | --- | --- |
| Complications | 41 | 37 |
| Genetics / Omics<br>(Genetics - 2009)<br>(Genetics/Genomes/Proteomics/Metabolomics - 2019) | 54 | 25 |
| Immunology and Transplantation | 24 | 11 |
| Islet studies | 37 | 25 |
| Obesity studies | 21 | 18 |
| Metabolism | 81 | 37 |
| Pathophysiology | 27 | 12 |
| New Methodology / Technology<br>(New Methodologies and Databases - 2009)<br>(Technological Advances - 2019) | 3 | 2 |
| Pharmacology | 8 | 9 |
| Signal Transduction | 16 | 2 |

### Funding status

We recorded the funding source for all papers in which this information was reported by the authors.

### Statistical analysis

All statistical analyses were performed using R^81^. For all analyses, generalized linear models were used. For mean score changes, a Poisson error distribution was assumed as scores are integers > −1. To calculate changes in proportions of scores, and the percentage of male- and female-biased studies, the scores were converted to a binary format and a binary (logistic) distribution was assumed. All analysis was started with a maximal model and *p*-values from the resulting ANOVA table are reported. *p* values are available in Supplemental File 3.

